# Cellular and molecular mechanisms involved in LTP induced by mild theta-burst stimulation in hippocampal slices from young male rats: from weaning to adulthood

**DOI:** 10.1101/2021.04.19.440440

**Authors:** NC Rodrigues, A Silva-Cruz, A Caulino-Rocha, A Bento-Oliveira, JA Ribeiro, D Cunha-Reis

## Abstract

Long-term potentiation (LTP) is a highly studied cellular process yet determining the transduction and GABAergic pathways that are the essential vs. modulatory for LTP elicited by theta-burst stimulation (TBS) in the hippocampal CA1 area is still elusive, due to the use of different TBS intensities, patterns, or different rodent/cellular models. We now characterized the developmental maturation and the transduction and GABAergic pathways required for mild TBS-induced LTP in hippocampal CA1 area in male rats. LTP induced by TBS (5×4) (5 bursts of 4 pulses delivered at 100Hz) lasted for up to 3h and was increasingly larger from weaning to adulthood. Stronger TBS patterns - TBS (15×4) or three TBS (15×4) separated by 6 min induced nearly maximal LTP not being the best choice to study the value of LTP-enhancing drugs. LTP induced by TBS (5×4) in young adults was fully dependent on NMDA receptor and CaMKII activity but independent of PKA or PKC activity. Furthermore, it was partially dependent on GABA_B_ receptor activation and was potentiated by GABA_A_ receptor blockade and less by GAT-1 transporter blockade. AMPA GluA1 phosphorylation on Ser_831_ (CaMKII target) but not GluA1 Ser_845_ (PKA target) was essential for LTP expression. The phosphorylation of the Kv4.2 channel was observed at Ser_438_ (CaMKII target) but not at Thr_602_ or Thr_607_ (ERK/MAPK pathway target). This suggests that cellular kinases like PKA, PKC or kinases of the ERK/MAPK family although important modulators of TBS (5×4)-induced LTP may not be essential for its expression in the CA1 area of the hippocampus.

**Graphical abstract:** 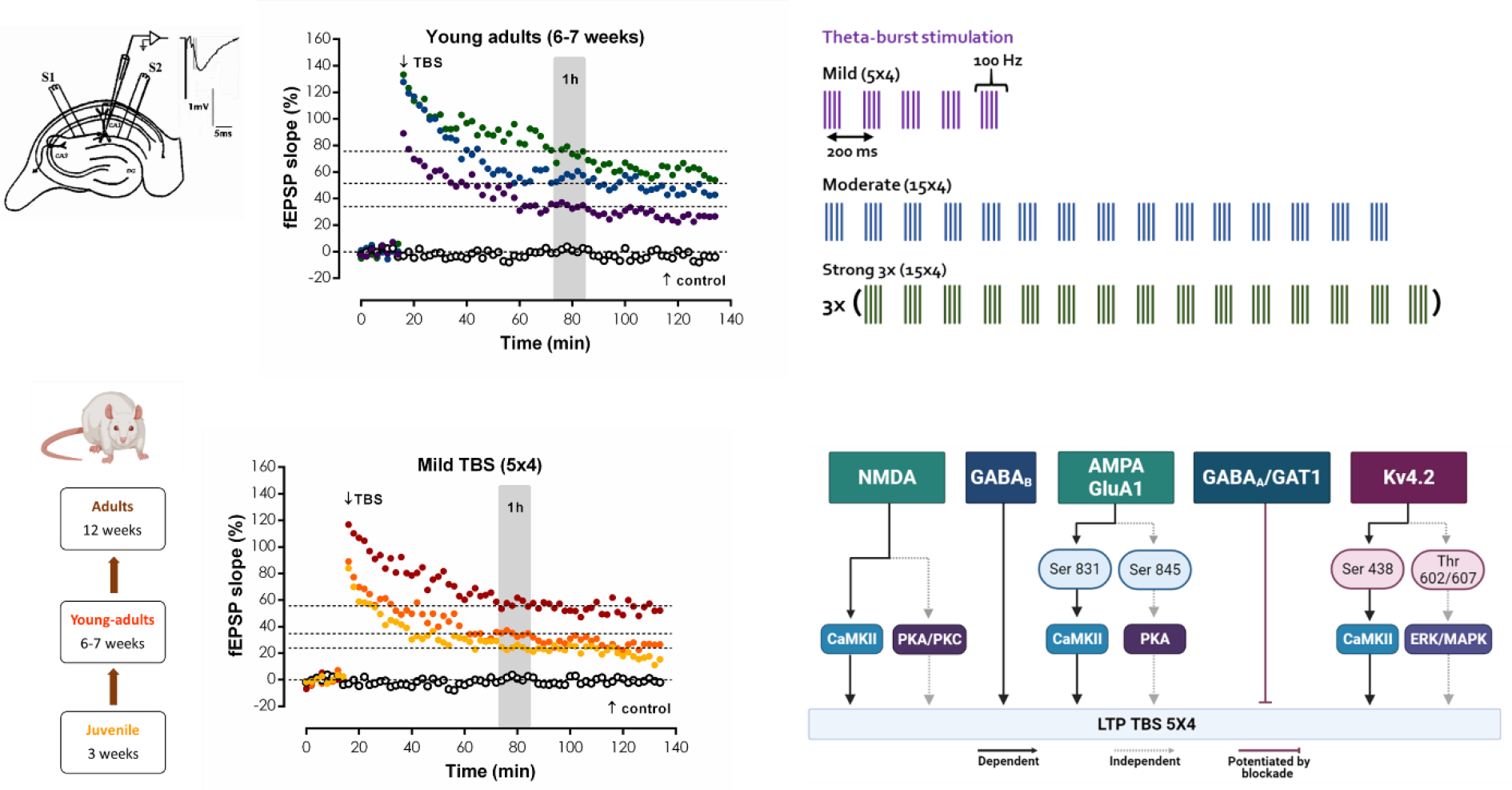

LTP induced by theta-burst stimulation (TBS) in young adult male rat hippocampal CA1 area is increasingly more robust with the number of bursts and number of spaced TBS trains. LTP induced by TBS shows developmental maturation increasing potentiation from weaning to adulthood. Expression of LTP induced by *mild TBS (5×4)* in young adults requires NMDA receptor and CaMKII activity and AMPA Ser831 phosphorylation, but not PKA or PKC activity, and is likely curtailed by Kv4.2 Ser438 phosphorylation.

## Introduction

Synaptic plasticity generated by activity dependent changes in the strength of synaptic communication, is widely accepted as the cellular mechanism underlying memory storage (Bliss and Collingridge, 2019). Long-term potentiation (LTP) evoked by high-frequency stimulation (HFS) was first described in 1973 in the contacts of perforant path fibres to granule cells in the dentate gyrus, and later in the remaining excitatory pathways of the hippocampus and was the first synaptic plasticity mode to be associated with hippocampus-dependent memory formation (Bliss and Collingridge, 1993; Lynch et al., 2007). The stability or long-lasting expression of LTP is dependent on the subsequent activation of multiple intracellular cascades that are differentially activated by various patterns of electrical activity *in vivo.* The magnitude and stability of LTP evaluated *in vitro* depends on several characteristics of the stimulus used to induce LTP such as frequency, intensity and pattern (Albensi et al., 2007; Baudry et al., 2015). Notwithstanding, intracellular cascades are differently activated in different rodent species and strains and at different animal developmental stages, making it difficult to set the stage for studies on the action of endogenous modulators and pharmacological tools on synaptic plasticity.

LTP can be associated to three mechanistically different processes (Park et al., 2018). Early-LTP can be triggered by a single episode of HFS, such as a tetanus or theta burst, requires activation of Ca^2+^/calmodulin-dependent protein kinase II (CaMKII) but is independent of both protein kinase A (PKA) and protein synthesis. Considerable evidence implicates an increase in the number of AMPA receptors (AMPARs) inserted into the postsynaptic membrane in early-LTP expression (Park et al., 2018). Late-LTP normally requires multiple episodes of HFS such as a tetani or theta burst for its induction; and these episodes need to be appropriately spaced in time (normally 10 min intervals). These protocols induce first an early-LTP, but then further mechanisms are activated, and the resulting potentiation thus involves both early and late-LTP. Late-LTP lasts more than 3h *in vitro*, is likely associated with synaptic contact enlargement and requires activation of PKA and *de novo* protein synthesis (Park et al., 2018). Longer-lasting forms of LTP require additional gene transcription and will not be addressed in this paper.

Theta burst stimulation (TBS) (Larson and Lynch, 1986; Rose and Dunwiddie, 1986) is a sequence of electrical stimuli that mimics CA3 and CA1 pyramidal complex-spike cell discharges observed during the hippocampal theta rhythm (3-7Hz). This EEG pattern has been associated with hippocampal memory storage and is believed to serve as a ‘tag’ for short term memory processing and to activate mechanisms eliciting early LTP (Vertes, 2005; Larson and Munkácsy, 2015). Bursts repeated at the theta frequency induce maximal LTP due to the suppression of feedforward inhibition by the first burst or a priming single pulse, that allows for enough depolarization to activate NMDA receptors (Larson and Lynch, 1988; Davies et al., 1990). This is mediated by activation of GABA_B_ autoreceptors that strongly inhibit GABA release from feedforward interneurons (Cobb et al., 1999) and synaptic GABA availability. For strong TBS trains the activation of presynaptic L-type Ca^2+^ channels also contributes to LTP induction (Morgan et al., 2001), but such intense stimulation patterns have been argued not to be required *in vivo* to induce maximal LTP (Larson and Munkácsy, 2015).

LTP induced by TBS is dependent on the elevation of dendritic Ca^2+^ resulting in the Ca^2+^-dependent activation and consequent auto-phosphorylation of CaMKII (Appleby et al., 2011; Kim et al., 2016). This results in the phosphorylation of AMPA receptor GluA1 subunits, that in turn promotes an enhancement of the channel conductance together with its recruitment to the active zone (Derkach et al., 1999; Appleby et al., 2011). It has been suggested that this recruitment may not always be required for the expression of hippocampal NMDA-dependent LTP (Henley and Wilkinson, 2016), and it is still controversial if it is essential to LTP stability (Henley and Wilkinson, 2016; Benke and Traynelis, 2019). Depending on the stimulation pattern and intensity used *in vitro* additional transduction pathways leading to the expression of late-LTP with involvement of PKA dependent phosphorylation of GluA1 subunits have been described (Huang Yan You and Kandel, 1994; Park et al., 2016) in the rat. Yet, LTP induced by a brief TBS in hippocampal slices elicits an LTP that is PKA-dependent in mice (Nguyen and Kandel, 1997). The diversity of rodent and cell culture models as well of TBS protocols described in the literature to elicit early and late LTP *in vitro* hinders the development of LTP modifying strategies targeting specific transduction pathways under firing conditions that are relevant for their activation *in vivo*.

The duration and extent of postsynaptic depolarization sustaining NMDA receptor activation, and ultimately leading to LTP induction, relies on the activity of both depolarizing and repolarizing postsynaptic currents and backpropagating action potentials. One channel that has been extensively studied regarding LTP expression is the Kv4.2 channel, that is largely responsible for the fast activating and deactivating A-current (I_A_) in CA1 pyramidal neuron distal dendrites, where it acts to control signal propagation and compartmentalization in dendrites (Beck and Yaari, 2008; Kim and Hoffman, 2008). Interestingly, it has also been shown that Kv4.2 expression is more prominent in dendritic spines than in dendritic shafts (Kim et al., 2007) and that LTP induction with strong TBS stimuli reduces the membrane levels of Kv4.2 on dendrites, which is accompanied by a shift in the voltage-dependence of I_A_, suggesting that the activity of the channel, dependent on modifications such as phosphorylation and/or trafficking, could contribute to LTP expression (Frick et al., 2004; Kim and Hoffman, 2008). Dendritic Ca^2+^ dynamics is also crucial to the activation of I_A_ during LTP induction, this being responsible for the precision of the time window allowed for LTP induction (Zhao et al., 2011). As above mentioned, the diversity of LTP induction protocols, cellular and animal models used to study the relevance of Kv4.2 activation and phosphorylation to the induction and expression of LTP complicates the evaluation of its actual contribution to LTP in different physiological conditions.

Although LTP induced by TBS is canonically related to GABA_B_-mediated suppression of feedforward inhibition and GABA_B_ receptor activation has been implicated in hippocampal-dependent learning and memory formation (Brucato et al., 1996), other GABAergic mechanisms and interneuron populations may contribute to LTP induced by mild TBS. Dendritic inhibition is ultimately mediated by synaptic GABA_A_ receptors that have distinct subunit composition from extra-synaptic GABA_A_ receptors (Farrant and Nusser, 2005; Brickley and Mody, 2012). The first mediate phasic inhibition, involved in the control of network activity, whereas extra-synaptic GABA_A_ receptors mediate tonic inhibition, occurring at a much slower time-window, shunting neuronal excitability and controlling neuronal input-output gain (Brickley and Mody, 2012). Thus, each GABA_A_ receptor population has likely a distinct role in regulation of TBS-induced LTP at neuronal dendrites. Synaptic inhibition at pyramidal cell dendrites has been demonstrated to be crucial for selective Ca^2+^-dependent input selectivity and precision of LTP induction (Müllner et al., 2015). Strong TBS has been demonstrated to induce long-term depression (LTD) in hippocampal interneurons *in vivo* and *in vitro* (Camiré and Topolnik, 2014; Lau et al., 2017), and recently mild TBS stimulation was used to induce both inhibitory LTD and inhibitory LTP at hippocampal GABAergic synapses (Udakis et al., 2020).

Mild TBS such as a single episode of five bursts (4 stimuli, 100Hz) delivered at 5Hz is a stimulation pattern that closely resembles the naturally occurring *in vivo* firing patterns in the hippocampal CA1 area during learning and memory acquisition and induces an early-LTP that is insensitive to PKA and protein synthesis inhibitors. This typically lasts for 2-3h *in vitro* in slices obtained from naïve rats where naturally occurring reinforcement or decay, either through behavioural experience or coincident motivational inputs are not present (Aidil-Carvalho et al., 2017; Çalışkan and Stork, 2018; Papaleonidopoulos and Papatheodoropoulos, 2018; Reyes-Garcia et al., 2018). Since the transduction pathways implicated in the expression of LTP induced by mild TBS in the rat hippocampus are unevenly described in the literature and depend often on the stimulation paradigm, animal model (and age) or in vitro preparation used, we re-evaluated the post-weaning to adulthood (3 to 12 weeks) expression of TBS-induced LTP, its dependence on stimulus strength and the involvement of different transduction pathways in LTP induced by mild TBS in rat hippocampal slices from young male rats (6-7 weeks).

## Material and Methods

The experiments were performed in hippocampal slices taken from juvenile (3 weeks old), young adult (6-7 weeks old) and adult (12 weeks old) male Wistar rats (Harlan Iberica, Barcelona, Spain) essentially as previously described (Cunha-Reis et al., 2014; Aidil-Carvalho et al., 2017) and were in agreement with the EU Directive 2010/63/EU for animal experiments. The animals were anesthetized with halothane, decapitated, and the right hippocampus dissected free in ice-cold artificial cerebrospinal fluid (aCSF) of the following composition in mM: NaCl 124, KCl 3, NaH_2_PO_4_ 1.25, NaHCO_3_ 26, MgSO_4_ 1.5, CaCl_2_ 2, glucose 10, and gassed with a 95% O_2_ - 5% CO_2_ mixture.

### LTP experiments

Hippocampal slices (400 µm thick) cut perpendicularly to the long axis of the hippocampus with a McIlwain tissue chopper were allowed energetic and functional recovery in a resting chamber in gassed aCSF at room temperature 22°C–25°C for at least 1 h. Each slice was transferred at a time to a submerged recording chamber of 1 ml capacity, where it was continuously superfused at a rate of 3 ml/min with the same gassed solution at 30.5°C. To obtain electrophysiological recordings slices were stimulated (rectangular pulses of 0.1 ms) through a bipolar concentric wire electrode placed on the Schaffer collateral/commissural fibres in the *stratum radiatum*. Two separate sets of the Schaffer pathway (S1 and S2) were stimulated Fig. 1A. Responses were evoked every 10 s alternately on the two pathways, each pathway being stimulated every 20 s (0.05Hz). The initial intensity of the stimulus was that eliciting a field excitatory post-synaptic potential (fEPSP) of 600–1000 mV amplitude, while minimizing contamination by the population spike, and of similar magnitude (about 50% of maximal slope) in both pathways. Evoked fEPSPs were recorded extracellularly from CA1 *stratum radiatum* (Fig. 1A) using micropipettes filled with 4 M NaCl and of 2–4 MΩ resistance. The averages of six consecutive responses from each pathway were obtained, measured, graphically plotted and recorded for further analysis with a personal computer using the LTP software (Anderson and Collingridge, 2001). The fEPSPs were quantified as the slope of the initial phase of the potential.

**Figure 1.**
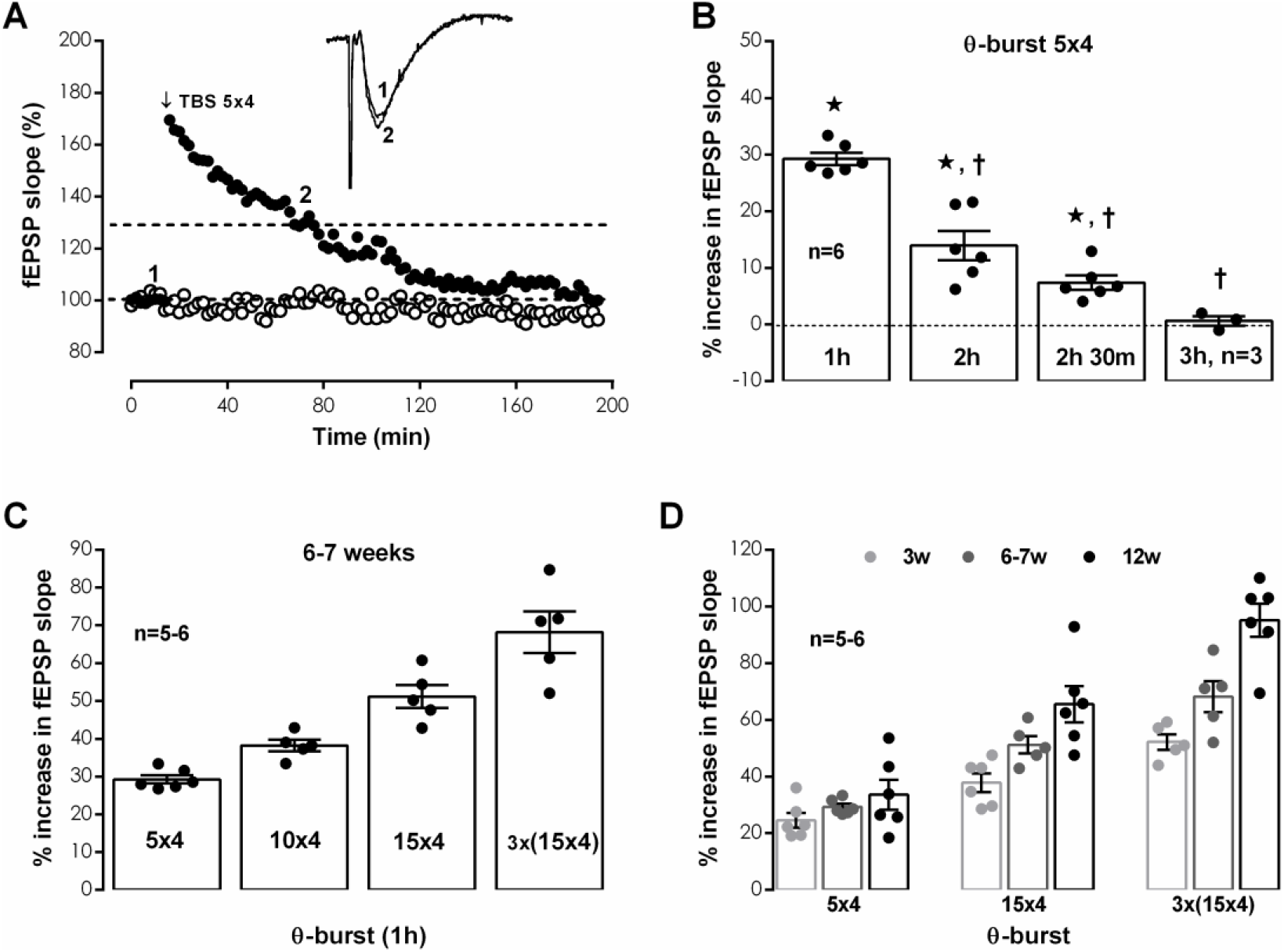
Hippocampal CA1 long-term potentiation of synaptic transmission elicited by different theta-burst stimulation protocols from weaning to adulthood. A. Time-course of changes in fEPSP slope caused by theta-burst stimulation (5 bursts at 5 Hz, each composed of four pulses at 100 Hz, *mild TBS(5×4)*) in a typical experiment for which a test pathway (-●-) was stimulated in the absence of added drugs and a control pathway (-○-) was used as internal control for TBS stimulation in the same hippocampal slice obtained from a young adult rat (6-7 weeks). *Inset:* Traces of fEPSPs obtained in the same experiment before (time point 1) and 50-60 min (time point 2) after theta burst stimulation. Traces are the average of eight consecutive responses and are composed of the stimulus artifact, the presynaptic volley and the fEPSP. B. LTP magnitude estimated from the averaged enhancement of fEPSP slope observed 50-60 min, 110-120 min, 140-150 min and 170-180 min after *mild TBS (5×4)* in the absence of added drugs in young adult rats (6-7 weeks). C. LTP magnitude obtained 50-60 min after theta-burst stimulation with increasingly stronger TBS stimulation paradigms: *mild TBS (5×4), TBS (10×4), moderate TBS (15×4)* and *strong TBS 3× (15×4)* bursts separated by 6 min in the absence of added drugs in young adult rats (6-7 weeks). D. Comparison of the LTP magnitude obtained 50-60 min after theta-burst stimulation with increasingly stronger TBS stimulation paradigms: *mild TBS (5×4), moderate TBS (15×4)* and *strong TBS 3× (15×4)* in *juvenile* (3 weeks), *young adult* (6-7 weeks) and *adult* (12 weeks) rats. Individual values and the mean ± S.E.M are depicted (B. - D.). *p < 0.05 (Student’s t test) as compared to the fEPSP slope before LTP induction; †p < 0.05 (one-way *ANOVA*, Tukey’s multiple comparison test) as compared with the potentiation obtained 50-60 min after *mild TBS (5×4)*.

The independence of the two pathways was tested at the end of the experiments by studying paired-pulse facilitation (PPF) across both pathways, less than 10% facilitation being usually observed. To elicit PPF, the two Schaffer pathways were stimulated with 50 ms interpulse interval. The synaptic facilitation was quantified as the ratio P2/P1 between the slopes of the fEPSP elicited by the second P2 and the first P1 stimuli.

LTP was induced by one of three growing intensity theta-burst stimulation (***TBS***) patterns: a ***mild TBS*** (five trains of 100 Hz, 4 stimuli, separated by 200 ms), a ***moderate TBS*** (fifteen trains of 100 Hz, 4 stimuli, separated by 200 ms) and a ***strong TBS*** stimulation pattern consisting of three moderate TBS stimulation trains delivered with a 6 min interval as previously used in our lab (Fontinha et al., 2008; Cunha-Reis et al., 2014; Aidil-Carvalho et al., 2017). The stimulation protocol used to induce LTP was only applied after having a stable fEPSP slope baseline for at least 20 min. The intensity of the stimulus was not changed during these stimulation protocols. LTP was quantified as the % change in the average slope of the potentials taken from 50 to 60 min after the induction protocol (except where otherwise stated), in relation to the average slope of the fEPSP measured during the 10 min that preceded the induction protocol. Control and test conditions (presence of drugs) were tested in independent slices. An independent pathway in the same slice was used as a control for TBS stimulation. In all experiments S1 always refers to the pathway (left or right, randomly assigned) to which TBS was applied. Test drugs were added to the perfusion solution 20 min before TBS stimulation and were present until the end of the experiment.

### Western blot analysis of GluA1 and Kv4.2 phosphorylation

For western blot studies, hippocampal slices were prepared as described above, allowed for functional recovery, and then placed in the electrophysiology chambers and superfused at a flow rate of 3 ml/min with gassed aCSF at 30.5 °C. Stimulation was delivered once every 15s in the form of rectangular pulses (0.1 ms duration) through a bipolar concentric wire electrode placed on the Schaffer collateral/commissural fibres in the *stratum radiatum* and lasted for 80 min (average duration of an electrophysiological experiment). In experimental conditions in which TBS was applied (but not in control chambers), TBS was delivered 20 min after the beginning of basal stimulation, and basal stimulation continued until the end of the experiment. For each experimental condition, hippocampal slices were collected in sucrose solution (320mM Sucrose, 1mg/ml BSA, 10mM HEPES e 1mM EDTA, pH 7,4) containing protease (complete, mini, EDTA-free Protease Inhibitor Cocktail, Sigma) and phosphatase (1 mM PMSF, 2 mM Na_3_VO_4_, and 10 mM NaF) inhibitors, homogenized with a Potter-Elvejham apparatus and centrifuged at 1500g for 10 min. The supernatant was collected and further centrifuged at 14000g for 12 min. The pellet was washed twice with modified aCSF (20mM HEPES, 1mM MgCl_2_, 1.2mM NaH_2_PO_4_, 2.7mM NaCl; 3mM KCl, 1.2mM CaCl_2_, 10mM glucose, pH 7.4) also containing protease and phosphatase inhibitors and resuspended in 300µl modified aCSF *per* hippocampus. Aliquots of this suspension of hippocampal membranes were snap-frozen in liquid nitrogen and stored at −80°C until use.

For western blot, samples incubated at 95°C for 5 min with Laemmli buffer (125mM Tris-BASE, 4% SDS, 50% glycerol, 0,02% Bromophenol Blue, 10% β-mercaptoethanol), were run on standard 10% sodium dodecyl sulphate polyacrylamide gel electrophoresis (SDS-PAGE) and transferred to PVDF membranes (Immobilon-P transfer membrane PVDF, pore size 0.45 μm, Immobilon). These were then blocked for 1 h with a 3% BSA solution, and incubated overnight at 4°C with rabbit antiphospho-Ser845-GluA1 (1:2000, Chemicon), rabbit antiphospho-Ser-831-GluA1 (1:3000, Chemicon), rabbit anti-GluA1 (1:4000, Millipore), rabbit anti-Kv4.2 (1:1000, Millipore), rabbit anti-phospho-Ser438-Kv4.2 (1:100, Santa Cruz Biotech), rabbit anti-phospho-Thr607-Kv4.2 (1:100, Santa Cruz Biotech), mouse monoclonal anti-phospho-Thr602-Kv4.2 (1:2000, Santa Cruz Biotech) and rabbit anti-alpha-tubulin (1:5000, Santa Cruz Biotech) primary antibodies or rabbit anti-beta-actin (1:10000, Proteintech) primary antibodies. After washing the membranes were incubated for 1h with anti-rabbit or anti-mouse IgG secondary antibody both conjugated with horseradish peroxidase (HRP) (Proteintech) at room temperature. HRP activity was visualized by enhanced chemiluminescence with Clarity ECL Western Blotting Detection System (Bio-Rad). Intensity of the bands was evaluated with the Image J software. Beta-actin density was used as a loading control. The % phosphorylation for each target on AMPA GluA1 subunits or Kv4.2 channels was determined by normalizing the band intensity of the phosphorylated form by the band intensity of the total GluA1 or Kv4.2.

### Statistics

Values are presented as the mean ± S.E.M of n experiments for electrophysiological studies and the mean ± S.E.M of n western blot experiments performed in duplicate for western-blot studies. A total of six animals per condition was used in all western blot experiments. For some of these western blot experiments slices from more than one animal were used per sample. Statistical analysis was performed using Prism software, GraphPad, San Diego, California. The significance of the differences between the means was calculated using the paired Student’s t-test when comparing two experimental groups, or with one-way analysis of variance ANOVA when comparing more than two experimental groups. P values of 0.05 or less were considered to represent statistically significant differences.

## Results

fEPSP obtained in hippocampal slices from *young-adult* (6-7 week-old) rats (Fig. 1.A, raw data from a single experiment) under basal stimulation conditions (40-60% of the maximal response in each slice) showed an average slope of 0.620±0.019 mV/ms (n=49). When *mild TBS* (5×4) was applied to the control pathway (S1) an LTP was observed, corresponding to a 29.6±1.2% increase in fEPSP slope (n=31, P<0.05) observed 50-60min after TBS. This potentiation progressively decayed until reaching average basal slopes 2h 30m to 3h after TBS 5×4 (n=3-6, Fig 1.B). Application of a second TBS train within 1h 20m to 2h latter in the test pathway (S2) in the absence of drugs always resulted in an LTP of similar magnitude (% increase in fEPSP slope of 32.4±3.6%, n=4) of the control pathway (S1), i.e., LTP obtained under these experimental conditions was similar in S1 and S2. Increasing the number of bursts to 10 or 15 (*moderate TBS*) enhanced the resulting potentiation evaluated 50-60min after TBS to a 38.2±1.5% (n=5, P<0.05) and 51.2±3.0% (n=5, P<0.05) enhancement in fEPSP slope (Fig 1.C), and also similar in S1 and S2. By applying a *strong TBS* paradigm (TBS 3× (15×4) separated by 6 min) potentiation evaluated 50-60min after stimulation was 68.2±5.5% (n=5, P<0.05, Fig 1.C).

In *juvenile* rats (3-4 weeks old) fEPSPs had an average slope of 0.596±0.026 mV/ms (n=22). Induction of LTP with a *mild TBS (5×4)* caused a smaller potentiation of the fEPSP slope observed 50-60min after stimulation (% increase in fEPSP slope 24.5±2.6%, n=6, P<0.05, Fig 1.D). *Moderate TBS (15×4)* and *strong TBS (3× TBS 15×4)* enhanced this potentiation to a 37.8±3.2% (n=6, P<0.05) and 52.2±2.7% (n=5, P<0.05) enhancement in fEPSP slope (Fig 1.D), respectively, that were also smaller than the one obtained in young adult rats for the same *TBS* paradigm. In *adult* rats (12-13 weeks old) the average fEPSP slope was of 0.635±0.022 mV/ms (n=32). The same *mild*, *moderate* and *strong TBS* paradigms caused a larger LTP and stronger enhancement in response to protocol strength than the ones observed in young adult rats. *Mild TBS* induced a potentiation of the fEPSP slope of 33.6±5.3%, n=6, P<0.05, Fig 1.D), while *moderate* and *strong TBS* enhanced this potentiation to a 65.5±6.4% (n=6, P<0.05) and 95.1±5.8% (n=5, P<0.05) enhancement in fEPSP slope (Fig 1.D) observed 50-60 min after stimulation, respectively. Except where otherwise stated, the involvement of the different transduction pathways in LTP induced by TBS was, as intended in this study, evaluated in young adult rats.

LTP expression induced by TBS in the hippocampus is believed to depend on suppression of feed-forward inhibition (Larson and Lynch, 1988), yet recent evidence strongly suggests that disinhibition, mediated by additional interneuron populations, is also crucial in the control of synaptic plasticity in the CA1 area of the hippocampus (Artinian and Lacaille, 2018; Udakis et al., 2020). To confirm the involvement of fast GABAergic transmission in hippocampal CA1 LTP induced by *mild TBS* we tested the influence of blocking GABA_A_ signaling using the selective GABA_A_ receptor antagonist bicuculline on LTP expression. When added to the slices bicuculline (10μM) increased fEPSP slope by 43.7±5.9% (n=5), also inducing an enhancement in dendritic excitability as reflected by an enhancement in the EPSP spike superimposition in the *stratum radiatum* of CA1 area. We thus reduced stimulation intensities to make fEPSP slopes of similar magnitude to the ones obtained in the absence of bicuculline. Stimulation with *mild TBS (5×4)* in the presence of bicuculine (10μM) caused a higher LTP than the one observed in control slices, corresponding to an enhancement of 44.0±4.7% (n=5, Fig. 2.A) of fEPSP slope observed 50-60 min after TBS. This difference in the response to *mild TBS (5×4)* was evident early after LTP induction since 5 min post stimulation the enhancement in fEPSP slope was significantly higher than the one observed in control slices. The resulting potentiation was also longer-lasting and showing no significant decay (P<0.05) when compared to the one observed in control conditions, as evidenced by the enhancement in fEPSP slope remaining 110-120min after TBS (29.5%±4.9%, n=5, in the presence of bicuculine vs. 13.9%±2.6%, n=6, in control conditions, Fig. 2.A, P<0.05). This suggests that additional inhibitory pathways mediate the influence of GABAergic transmission on TBS-induced LTP induction and maintenance. GABA_B_ autoreceptor activation by enhanced GABA release has been implicated in the suppression of feedforward inhibition during TBS (Brucato et al., 1996). Synaptically released GABA is subject to reuptake by high-affinity Na^+^- and Cl^-^-dependent GABA transporters (GATs) (Cunha-Reis et al., 2017). GAT1 is predominantly expressed in presynaptic GABAergic boutons and plays a crucial role in controlling GABA spillover and, consequently, modulation of both phasic and tonic GABAergic inhibition (Gong et al., 2009). As such, we tested the role of synaptic GABA availability in the induction and expression of LTP induced by *mild TBS* upon partial inhibition of GAT1 transporters. When added to the slices, the selective GAT1 antagonist SKF89976a (5μM) did not significantly change fEPSP slope. Unlike what was previously observed in mice (Gong et al., 2009), upon blockade of GAT1 transporters with SKF89976a (5μM), LTP induction with *mild TBS (5×4)* caused a long-lasting enhancement of 40.6±2.9% (n=6, Fig. 2.B) in fEPSP slope 50-60 min after stimulation, that was higher (P<0.05) than the one observed in control conditions 29.7±3.4% (n=6), but this difference was attenuated 110-120min after stimulation (Fig. 2.B). Finally, we tested the influence of GABA_B_ receptor blockade with the selective antagonist CGP55845 on the induction and expression of LTP induced by *mild TBS (5×4)*. When CGP55845 (1μM) was present in the bath, stimulation with *mild TBS (5×4)* elicited a weaker LTP than the one observed in control slices, corresponding to an enhancement of fEPSP slope of 17.8±0.5% (n=5, Fig. 2.C) of fEPSP slope observed 50-60 min after TBS, and that was nearly extinct (% increase in fEPSP slope: 4.7±1.5%, n=5) 110-120 min after TBS, confirming the importance of activation of GABA_B_ receptors to the induction of LTP by TBS.

**Figure 2.**
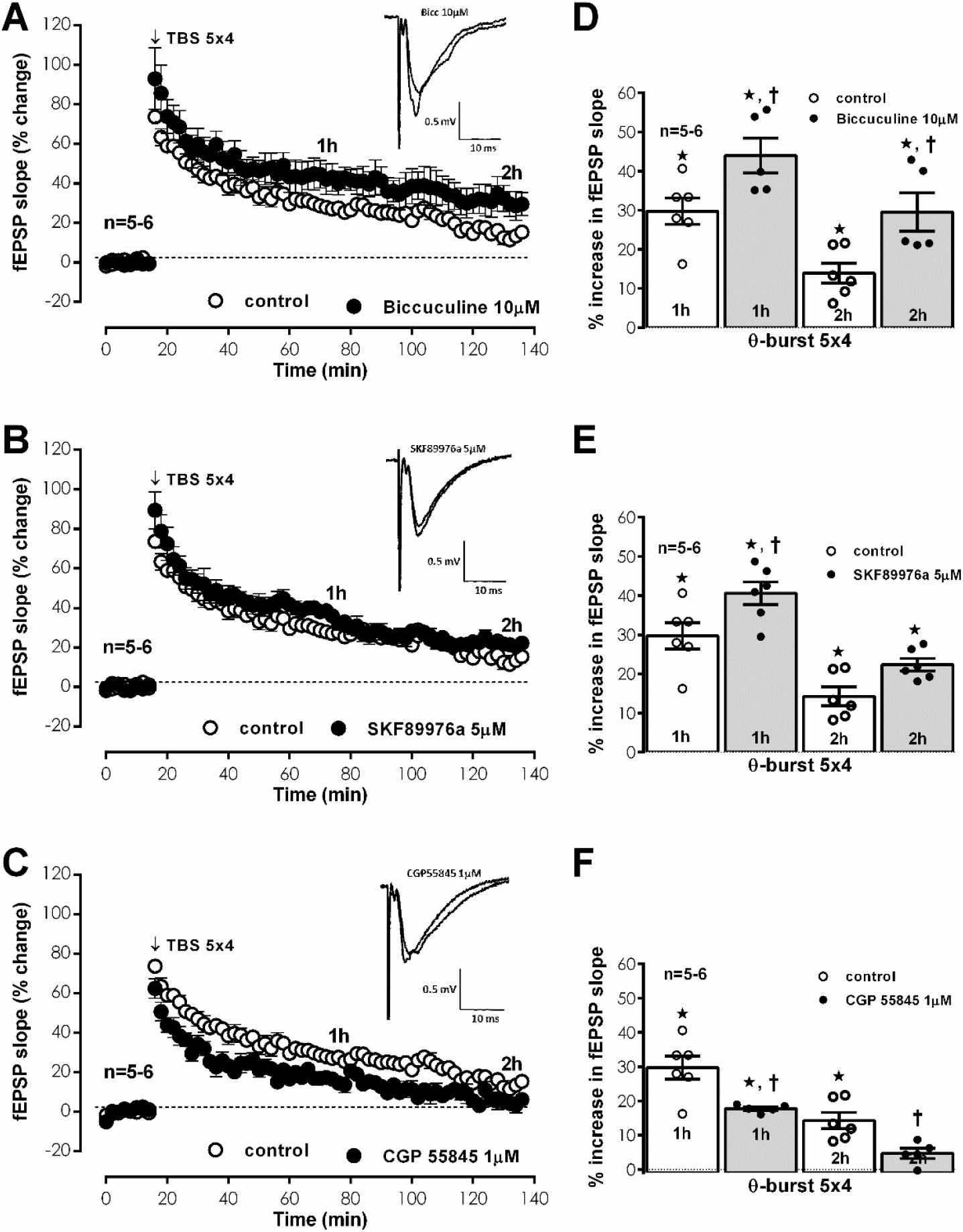
Hippocampal CA1 LTP induced by *mild TBS (5×4)* is dependent on GABAergic transmission. **A., B.** and **C.** Averaged time-course of changes in fEPSP slope caused by theta-burst stimulation (*mild TBS (5×4))* in the absence (-○-) and in the presence (-●-) of the selective GABA_A_ receptor antagonist Bicuculline (10μM, **A.**), the selective GAT-1 GABA transporter inhibitor SKF8997A (5μM, **B.**) or the selective GABA_B_ receptor antagonist CGP 55845 (1μM, **C.**). Control and test conditions (absence and presence added drugs) were tested in independent slices. ***Inset:*** Traces of fEPSPs obtained in the same experiment before (0-16min) and 50-60 min after (**1h**) *mild TBS (5×4)* stimulation in the presence of the respective drugs. Traces are the average of eight consecutive responses and are composed of the stimulus artifact, the presynaptic volley and the fEPSP. **D., E.** and **F.** Magnitude of LTP estimated from the averaged enhancement of fEPSP slope observed 50-60 and 110-120 min after *mild TBS (5×4)* in the absence (-○-) or in the presence (-●-) the selective GABA_A_ receptor antagonist Bicuculline (10μM, **A.**), the selective GAT-1 GABA transporter inhibitor SKF8997A (5μM, **B.**) or the selective GABA_B_ receptor antagonist CGP 55845 (1μM, **C.**). Individual values and the mean ± S.E.M are depicted (**D.** - **F.**). *p < 0.05 (Student’s t test) as compared to the fEPSP slope before LTP induction; †p < 0.05 (one-way *ANOVA*, Tukey’s multiple comparison test) as compared with the potentiation obtained in different conditions and time points in the same slices.

To investigate if LTP induced by *mild TBS* was dependent on the activation of NMDA receptors we tested the effect *mild TBS (5×4)* in the presence of the NMDA receptor antagonist (2R)-amino-5-phosphonopentanoate (AP-5). As expected under basal stimulation conditions, when added to the slices from the beginning of experiments AP-5 (100μM) did not significantly change fEPSP slope. In the presence of AP-5 (100μM), stimulation with *mild TBS (5×4)* did not elicit an LTP (P>0.05, n=4, Fig. 3.A), as previously described in the same experimental conditions (e. g. Aidil-Carvalho et al., 2017).

**Figure 3.**
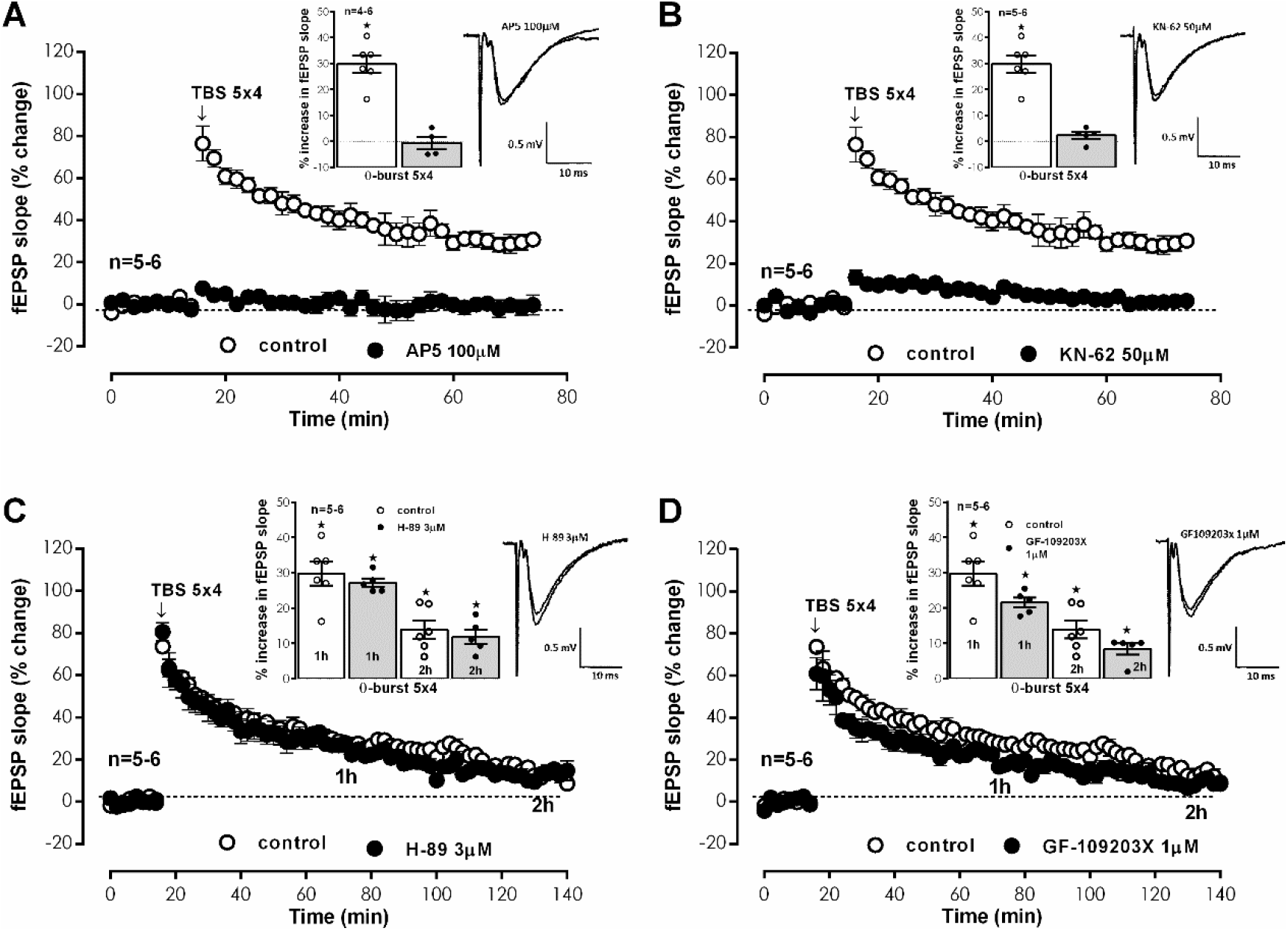
Hippocampal CA1 LTP induced by *mild TBS (5×4)* in young adult rats is dependent on NMDA activation and CaMKII activity but not PKA or PKC activity. **A.-D.** Averaged time-course of changes in fEPSP slope induced by *mild TBS (5×4)* in the absence (-○-) and in the presence (-●-) of either the selective NMDA receptor antagonist AP5 (100μM, **A.**), the selective CaMKII inhibitor KN-62 (50μM, **B.**), the selective PKA inhibitor H-89 (3μM, **C.**) or the selective PKC inhibitor GF109203x (1μM, **D.**). Control and test conditions (absence and presence of added drugs) were tested in independent slices. *Inset:* Traces of fEPSPs (right) obtained in the same experiment before (0-16min) and 50-60 min after (**1h**) *mild TBS (5×4)* stimulation and magnitude of LTP (left) estimated from the averaged enhancement of fEPSP slope observed 50-60 min after theta-burst stimulation in the absence (-○-) or in the presence (- ●-) of the NMDA receptor antagonist AP5 (100μM, **A.**), the selective CaMKII inhibitor KN-62 (50μM, **B.**), the selective PKA inhibitor H-89 (3μM, **C.**) or the selective PKC inhibitor GF109203x (1μM, **D.**). Individual values and the mean ± S.E.M are depicted (**A.** - **D.**). *p < 0.05 (Student’s t test) as compared to the fEPSP slope before LTP induction; †p < 0.05 (one-way *ANOVA*, Tukey’s multiple comparison test) as compared with the potentiation obtained in control conditions for the same time point in the same slices.

It is generally accepted the expression of LTP relies on the Ca^2+^-dependent activation and consequent auto-phosphorylation of CaMKII, that then elicits the recruitment of AMPA GluA1 subunits (Appleby et al., 2011; Park et al., 2021). However, it has been reported that this is not required for the expression of NMDA-dependent LTP in different hippocampal subfields in the mouse (Cooke et al., 2006). Involvement of CaMKII in LTP induced by *mild TBS (5×4)* was investigated using the CaMKII selective inhibitor KN-62. When added to the slices KN-62 (50μM) did not significantly change fEPSP slope. When *mild TBS (5×4)* was delivered in the presence of KN-62 (50μM) LTP expression was suppressed (% change in fEPSP slope of 2.2±1.4%, n=4, Fig. 3.B). These results confirm that LTP induced by *mild TBS (5×4)* is fully dependent on CamKII activity. We further investigated the involvement of the G_s_/adenylate cyclase/PKA transduction system in CA1 on LTP induced by *mild TBS (5×4)*. In the presence of the selective PKA inhibitor (Chijiwa et al., 1990) H-89 (3μM), LTP induced by *mild TBS (5×4)* was not changed (% increase in fEPSP slope 50-60 min after stimulation: 27.2±1.2%, n=5, Fig. 3.C, P<0.05). In view of the absence of effect of the selective PKA inhibitor LTP induced by *mild TBS (5×4)*, the involvement of PKC was also investigated. Upon selective inhibition of PKC with GF 109203X (1 μM) (Toullec et al., 1991), *mild TBS (5×4)* elicited an LTP that showed similarly little change (% increase in fEPSP slope 50-60 min after stimulation: 21.5±1.5%, n=5, Fig. 3.D, P>0.05). Expression of late-LTP induced by *moderate TBS (15×4)* in C57BL/6J mice is dependent on PKA activity and protein synthesis (Nguyen and Kandel, 1997), yet only multiple spaced TBS trains have been described to induce an LTP dependent on PKA in hippocampal slices from Sprague-Dawley rats (Park et al., 2016, 2021). To confirm this, we readdressed the PKA-dependency of TBS induced LTP by using a *moderate TBS (15×4)*, as used by Nguyen and Kandel in 1997 and a *strong TBS 3×(15×4)* separated by 6 min, that would assure the conditions of three TBS bursts with sufficient spacing to mimic the spaced TBS used by Park et al. (2016), and in rats of similar age to the ones in the mouse study (12 weeks). Upon *moderate TBS (15×4)*, the resulting potentiation in fEPSP slope evaluated 50-60min after TBS was 65.5±6.4% (n=6, P<0.05, Fig 4.A) and 110-120 min after stimulation showed very little decay 61.2±9.3% (n=6, P<0.05, Fig 4.A). In the presence of the selective PKA inhibitor H-89 (3μM), the potentiation of fEPSP slope induced by *moderate TBS (15×4)* was not significantly changed either 50-60 min post stimulation (62.5±2.1%, n=5, Fig. 4.A) or 110-120 min post stimulation (63.4±4.2%, n=5, Fig. 4.A).

**Figure 4.**
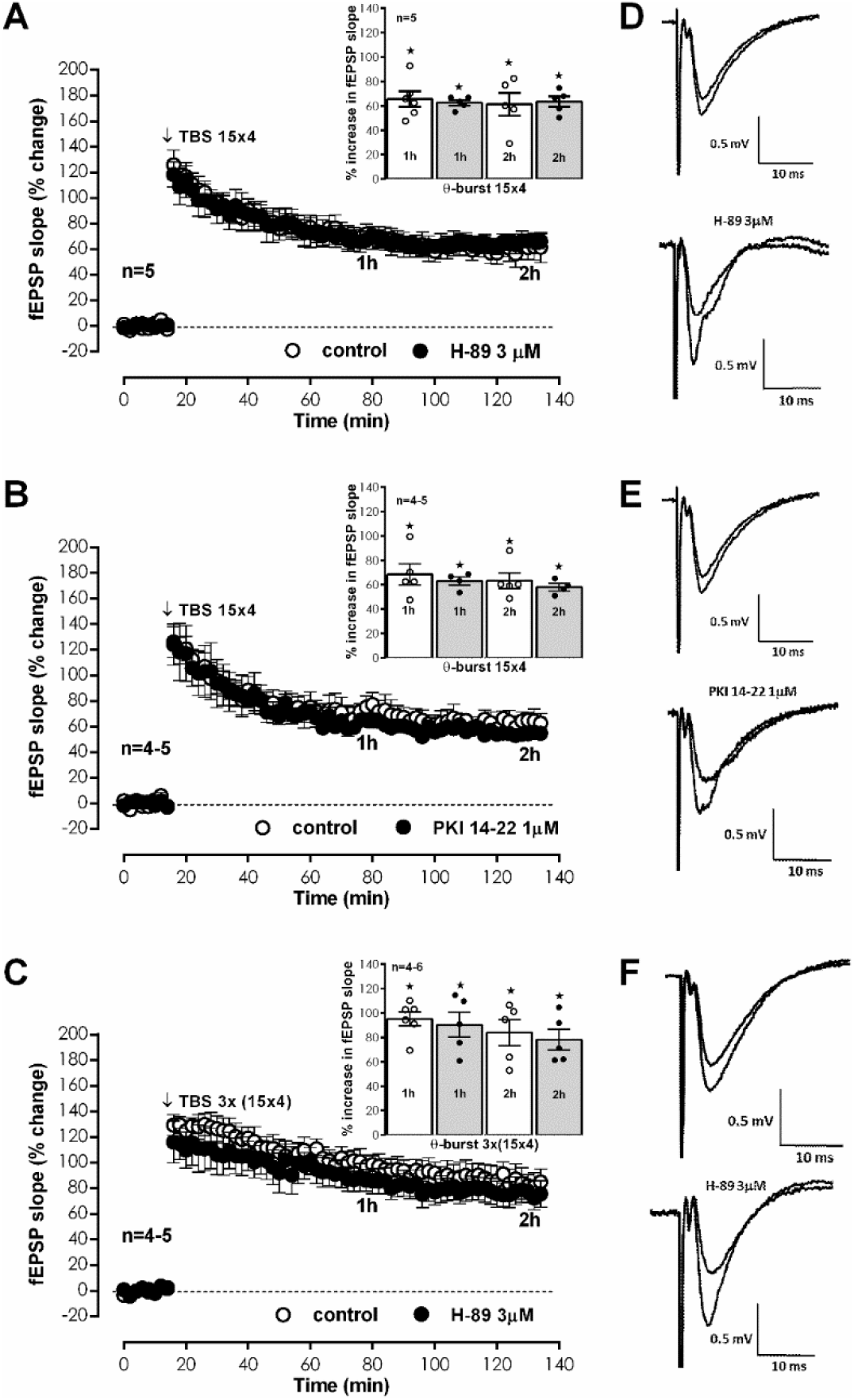
Hippocampal CA1 LTP induced by *moderate TBS (15×4)* and *strong TBS 3× (15×4)* in adult rats is independent of PKA activity. **A.-B.** Averaged time-course of changes in fEPSP slope induced by *moderate TBS (15×4)* in the absence (-○-) and in the presence (-●-) of either the selective PKA inhibitor H-89 (3μM, **A.**) or another selective PKA inhibitor PKI 14-22 (1μM, **B.**). Control and test conditions (absence and presence of added drugs) were tested in independent pathways in the same slice obtained from adult rats. *Inset:* Magnitude of LTP estimated from the averaged enhancement of fEPSP slope observed 50-60 or 110-120 min after *moderate TBS (15×4)* in the absence (-○-) and in the presence (-●-) of either the selective PKA inhibitor H-89 (3μM, **A.**) or the selective PKA inhibitor PKI 14-22 (1μM, **B.**).**C.** Averaged time-course of changes in fEPSP slope induced by *strong TBS 3× (15×4)* in the absence (-○-) and in the presence (-●-) of the selective PKA inhibitor H-89 (3μM) *Inset:* Magnitude of LTP estimated from the averaged enhancement of fEPSP slope observed 50-60 or 110-120 min after *strong TBS 3× (15×4)* in the absence (-○-) and in the presence (-●-) of the selective PKA inhibitor H-89 (3μM). Individual values and the mean ± S.E.M are depicted (**A.** - **C.**). **D. - F.** Traces of fEPSPs) obtained in the same experiment before (0-16min) and 50-60 min after (**1h**) either *moderate TBS (15×4)* (**D. – E.**) or strong *strong TBS 3× (15×4)* (**F.**) stimulation in the absence or in the presence (-●-) of the selective PKA inhibitor H-89 (3μM, **D., F.**) or the selective PKA inhibitor PKI 14-22 (1μM, **E.**). *p < 0.05 (Student’s t test) as compared to the fEPSP slope before LTP induction; †p < 0.05 (one-way *ANOVA*, Tukey’s multiple comparison test) as compared with the potentiation obtained in control conditions for the same time point in the same slices.

H-89 is a competitive antagonist at the ATP binding site of the PKA catalytic subunit and its EC_50_ is known to depend strongly on intracellular ATP concentrations. Since the absence of effect of PKA on late-LTP could be due its ineffective inhibition of PKA, we used another inhibitor, PKI 14-22 amide, a PKA inhibitor that binds with high-affinity the catalytic subunit of PKA, similarly to the binding of the PKA regulatory subunit, and that promotes inhibition of the kinase activity a K_i_ in the nanomolar range (Glass et al., 1989). The presence of the selective PKA inhibitor PKI 14-22 amide (1μM), again did not significantly change (P>0.05) the potentiation of fEPSP slope induced by *moderate TBS (15×4)* either 50-60 min post stimulation (63.0±3.4%, n=4, Fig. 4.B) or 110-120 min post stimulation (58.0±3.0%, n=4, Fig. 4.B).

Upon *strong TBS 3× (15×4)*, the resulting potentiation in fEPSP slope evaluated 50-60min after TBS was 95.1±5.8% (n=6, P<0.05, Fig. 4.C) and showed only a small decay since the enhancement in fEPSP slope observed 110-120 min after stimulation was 83.8±10.7% (n=6, P<0.05, Fig. 4.C). Under these experimental conditions the selective PKA inhibitor H-89 (3μM) also did not significantly change (P>0.05) the potentiation of fEPSP slope induced by *strong TBS 3× (15×4)* either 50-60 min post stimulation (90.3±10.1%, n=5, Fig. 4.C) or 110-120 min post stimulation (78.2±8.6%, n=5, Fig. 4.C).

To further investigate the transduction pathways involved in the expression of LTP induced *mild TBS (5×4),* we investigated the downstream targets of different intracellular kinases on AMPA GluA1 receptor subunits, previously reported to mediate expression of hippocampal LTP by promoting traffic or modifying opening probability of AMPA receptors (Lee et al., 2003). When GluA1 phosphorylation was inspected 50 min after *mild TBS (5×4),* no significant changes in Ser_845_ phosphorylation (a site targeted by PKA) or in the total expression of GluA1 subunits (P>0.05, n=3-4, Fig 5A-B, E-F) were observed. Yet, an enhancement of 62.1±11.0%, n=4 (Fig. 5.C and D) in GluA1 phosphorylation in Ser_831_ residue, a phosphorylation site targeted by both CaMKII and PKC, was observed under these experimental conditions. Increasing the TBS stimulus to a *moderate TBS (15×4)* did not change the pattern of GluA1 phosphorylation but increased, although not significantly when compared to *TBS(5×4)* (P>0.05), the level of GluA1 phosphorylation at Ser_831_ by 91.3±10.4%, n=3 (Fig. 5.C and D).

**Figure 5.**
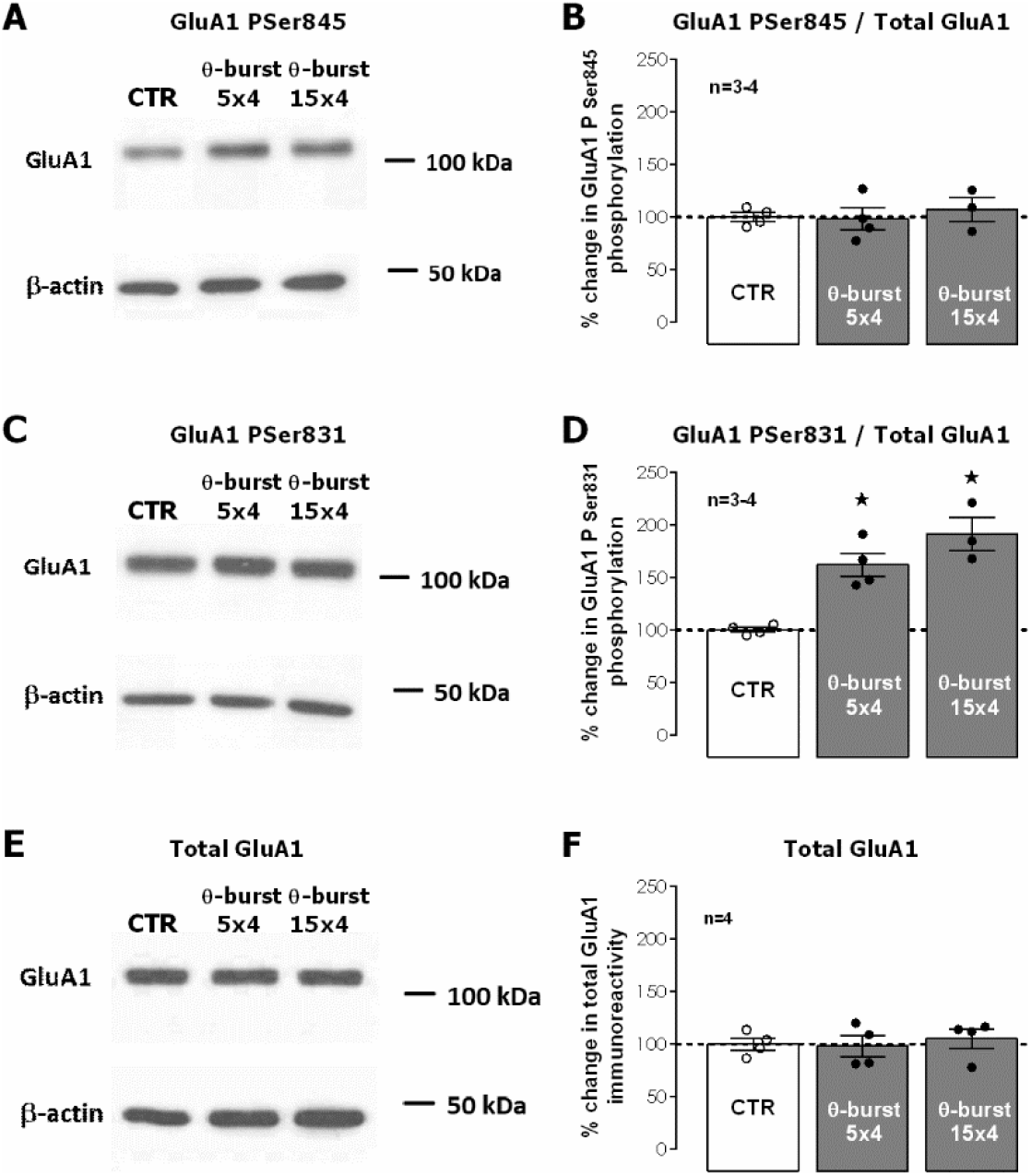
Impact of theta-burst stimulation on phosphorylation of hippocampal AMPA GluA1 subunits on Ser_845_ and Ser_831_. **A., C.** and **E.** Western-blot immunodetection of AMPA GluA1 phosphorylated forms in Ser_845_ and Ser_831_ and of total GluA1 subunits obtained in one individual experiment in which hippocampal slices were subjected to Schaffer collateral basal, *mild TBS (5×4)* or *moderate TBS (15×4)* stimulation. Slices were monitored for 50 min after TBS (or equivalent time for controls) before WB analysis. **B.** Total GluA1 immunoreactivity and **D.** and **F.** % GluA1 phosphorylation on Ser_845_ and Ser_831_ residues of GluA1 subunits. Individual values and the mean ± S.E.M of four independent experiments performed in duplicate are depicted (**B., D.,** and **F**). 100% - averaged GluA1 immunoreactivity or GluA1 phosphorylation obtained in control conditions (absence of TBS). **\*** represents p < 0.05 (*ANOVA*, Tukey’s multiple comparison test) as compared to absence of TBS.

Finally, we investigated the downstream targets of different intracellular kinases on Kv4.2 dendritic K^+^ channels, previously reported to contribute to the expression of hippocampal LTP by suppressing the A-current and facilitating action potential backpropagation (Frick et al., 2004; Rosenkranz et al., 2009). When Kv4.2 phosphorylation was inspected 50 min after *mild TBS (5×4),* a significant enhancement (119.8±43.8%, P<0.05, n=4, Fig 6.A-B) was observed in Ser_438_ phosphorylation (a site targeted by CaMKII) but not in the total expression of Kv4.2 subunits (P>0.05, n=4, Fig 6.C-D). No significant changes (P>0.05, n=3) were observed in Kv4.2 phosphorylation in two ERK (extracellular signal-regulated kinases) phosphorylation sites Thr_607_ (Fig 6.E) and Thr_602_ (Fig 6.F) and basal phosphorylation (control conditions) at these two sites was low.

**Figure 6.**
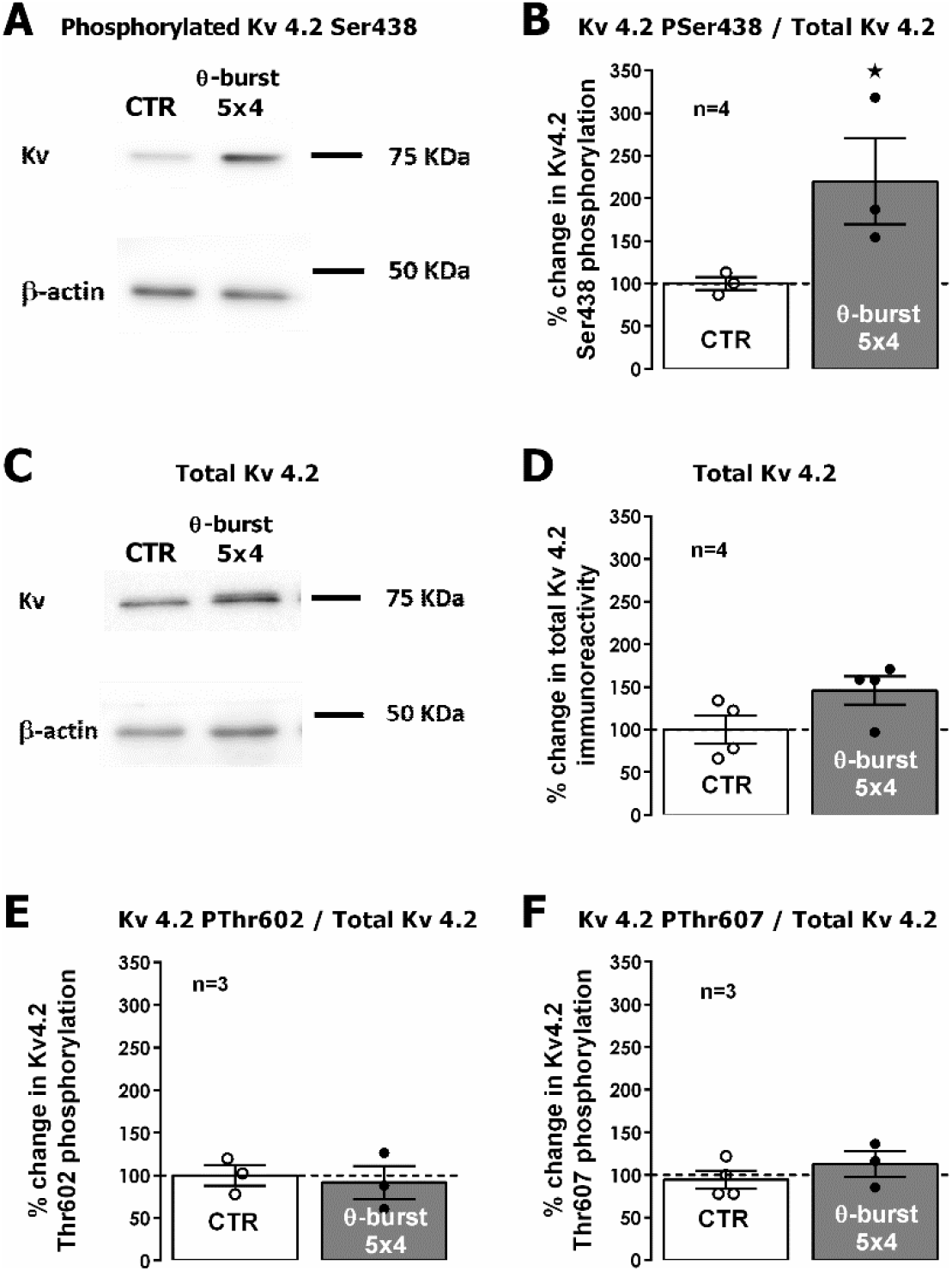
Impact of theta-burst stimulation on phosphorylation of hippocampal Kv4.2 on Ser_438_, Thr_602_ and Thr _607_. **A.** and **C.** Western blot immunodetection of Kv4.2 phosphorylated forms in Ser438 and total Kv4.2 obtained in one individual experiment in which hippocampal slices were subjected to Schaffer collateral basal or *mild TBS (5×4)* stimulation. Slices were monitored for 50 min after TBS (or equivalent time for controls) before WB analysis. **B., D., E,** and **F.** Averaged total Kv4.2 immunoreactivity and immunoreactivity of its phosphorylated forms in Ser_438_, Thr_602_ or Thr_607_, normalized to the total variation in Kv4.2. Individual values and the mean ± S.E.M of 3-4 independent experiments performed in duplicate are depicted (**B., D., E,** and **F**). 100% - averaged Kv4.2 immunoreactivity obtained in control conditions (absence of TBS). **\*** represents p < 0.05 (*Student’s t* test) as compared to absence of TBS.

## Discussion

The main findings of the present work are that: 1) *mild TBS* stimulation (5 x 4 (100Hz) stimuli, separated by 200ms), a stimulation paradigm mimicking the duration and pattern of complex-spike discharges under theta-rhythm modulation that are observed during exploration and memory acquisition, induces a long-lasting potentiation of synaptic transmission that is increasingly larger from weaning (3 week-old) through adulthood (12-week-old) in the CA1 area of hippocampal slices obtained from male Wistar rats; 2) Increasing the duration of the train to 15 x 4 (100Hz) stimuli, separated by 200ms (*moderate TBS*) or concomitantly increasing the duration and number of these trains to three (*strong TBS*) resulted in an increased potentiation and stability of the elicited LTP at the three developmental stages tested (3-, 7- and 12 week-old); 3) The LTP elicited by *mild TBS* in young (7-week-old) rats was dependent on GABA_B_ receptor activation and is endogenously inhibited by GABA_A_ receptor activation and by GAT-1 activity; 4) *mild TBS* induced LTP in young rats was fully dependent on NMDA receptor and CaMKII activity and for young adult to adult rats and the two stronger TBS paradigms was independent of PKA activity; 5) Furthermore, LTP induced by *mild TBS* in young rats results in putative CaMKII-mediated phosphorylation of GluA1 AMPA receptor subunits at Ser831 and of Kv4.2 K^+^ channels at Ser438 (Fig. 7).

**Figure 7.**
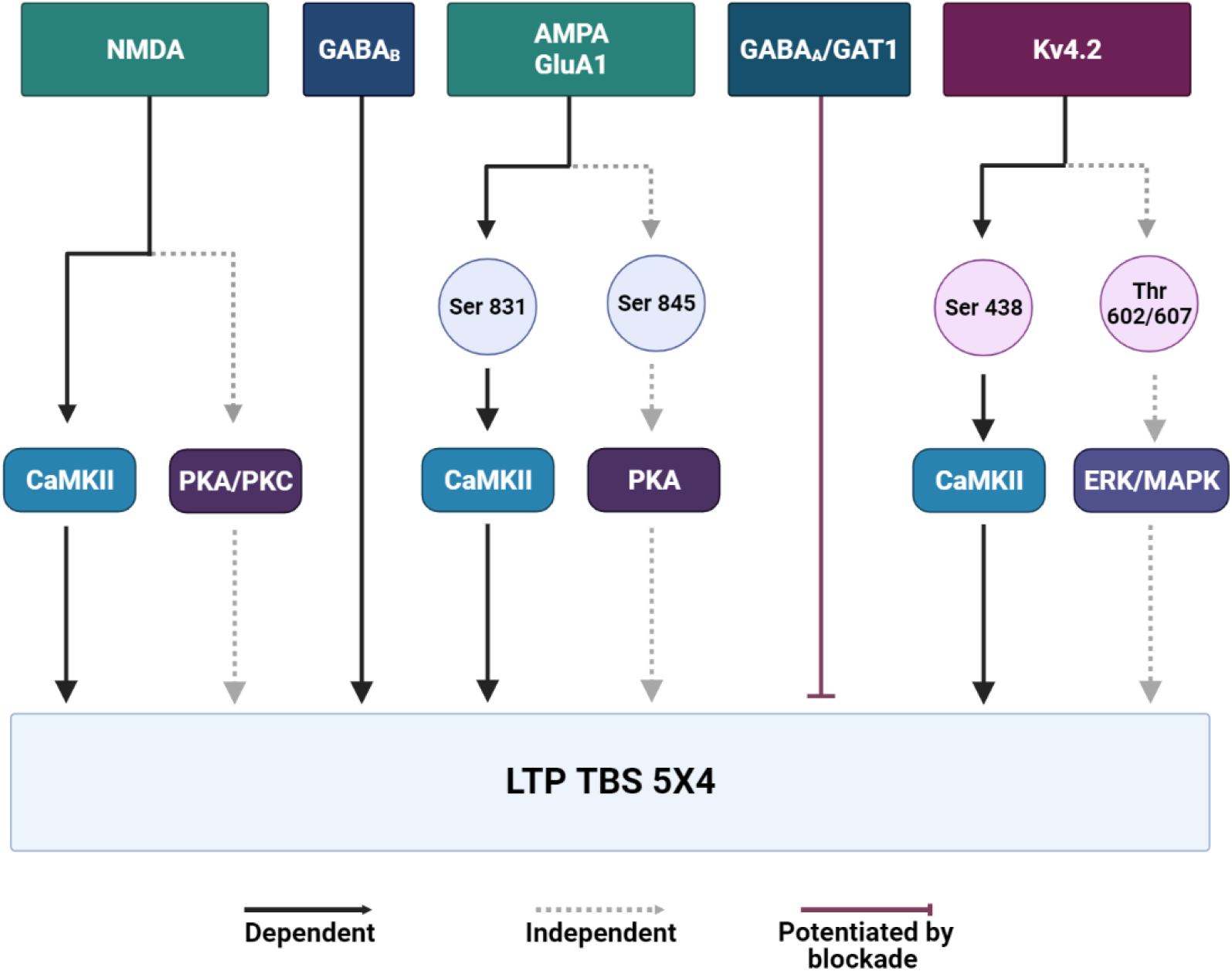
Schematic representation of the main transduction pathways required for the expression of LTP in the CA1 area of the hippocampus induced by different TBS intensities in young adult rats. AMPA - α-amino-3-hydroxy-5-methyl-4-isoxazolepropionic acid ionotropic receptor for glutamate; CaMKII – Ca2+/calmodulin-dependent protein kinase II; ERK - extracellular signal-regulated kinase; GABA_A_ – receptor A for GABA (ionotropic); GABA_B_-receptor B for GABA (metabotropic); GAT-1 – GABA transporter 1; Kv4.2 – voltage-gated potassium channel 4.2; MAPK - mitogen-activated protein kinases; NMDA –N-methyl D-aspartate ionotropic receptor for glutamate; PKA – protein kinase A; PKC – protein kinase C

In the present study *mild TBS* (5 bursts, 4 pulses delivered at 100Hz), elicited a long-lasting potentiation of fEPSP slope in the CA1 area of male rat hippocampal slices that lasted at least 2h. This potentiation was increasingly larger from 3-, 7-, to 12-week-old rats. Previous studies showed that during postnatal development there is a reinforcement of the cellular mechanisms leading to LTP expression and stability (Kramár and Lynch, 2003; Cao and Harris, 2012; Larson and Munkácsy, 2015) but it was never unequivocally shown in the same study that this enhancement in potentiation continues during postweaning development until adulthood (12-week-old rats). In fact, some studies study indifferently animals aged 4-12 weeks (Park et al., 2016, 2021), and thus do not discriminate differences in LTP during post-weaning development to adulthood. Stronger TBS patterns, achieved by increasing the number of bursts to 10-15 (*moderate TBS*) or delivery of three of these trains separated by 6 min (*strong TBS*), further enhanced the resulting potentiation for all age groups studied. *Mild TBS* constitutes an ideal intensity of TBS for pharmacological studies aiming to improve LTP outcome, since increasing the number of pulses as in *moderate TBS* (e.g. 15 bursts, 4 pulses) elicits a nearly maximal LTP for this stimulation pattern. This ceiling effect could only be overcome by delivering several TBS trains as previously described (Larson and Munkácsy, 2015). Weaker TBS stimulation (3 bursts, 4 pulses) was previously observed not to elicit a long-lasting potentiation of fEPSP slope at the developmental ages tested in this study (Costenla et al., 1999). Also, as previously reported (Aidil-Carvalho et al., 2017), the potentiation elicited by *mild TBS* in the CA1 area of the rat hippocampus was fully dependent on NMDA receptor activity (Fig. 3.A), unlike the potentiation elicited by other stronger and slightly differently patterned TBS (200Hz tetanus frequency) for which a concomitant dependency on voltage-gated calcium channels (VGCCs) was observed (Grover and Teyler, 1990; Grover, 1998).

Suppression of feed-forward phasic inhibition, mediated by GABA_B_ autoreceptor suppression of GABA release, is believed to mediate induction of LTP by TBS in excitatory synapses of the hippocampus (Pacelli et al., 1989; Grover and Yan, 1999; Larson and Munkácsy, 2015), allowing for enough temporal summation of excitation and prolonged depolarization that ultimately lead to NMDA receptor activation. In addition, *mild* TBS also activates cooperative postsynaptic mechanisms dependent on GABA_B_ and metabotropic glutamate receptors, leading to a sustained potentiation of fast GABA_A_ synaptic transmission in GABAergic synapses to pyramidal cells (Patenaude et al., 2003). Despite this, the net result is a potentiation of synaptic transmission as evaluated by the long-lasting changes in fEPSP slope. In this study, blocking GABA_A_ signaling using supramaximal concentrations of the selective GABA_A_ receptor antagonist bicuculline resulted in an enhancement of the magnitude of LTP induced by *mild* TBS in the CA1 area. This is consistent with previous studies in Sprague-Dawley rats using primed-burst protocols (Pacelli et al., 1989) and much stronger TBS protocols using either 100Hz or 200Hz tetanus frequency (Chapman et al., 1998; Grover and Yan, 1999). The resulting potentiation lasted longer than the one observed in control conditions and showed little decay 2h after induction, unlike the one obtained in control conditions that was run down to one third of the potentiation observed 1h after induction. Since sustained changes in GABA_A_ mediated currents in response to *mild* TBS do not contribute to the fEPSP slope in the presence of bicuculine, this result likely reflects the potentiation (and its stability) elicited by TBS at glutamatergic synapses and suggests that additional inhibitory pathways (other than suppression of phasic feed-forward inhibition during induction) mediate the influence of GABAergic transmission on the expression and maintenance of TBS-induced LTP. One hypothesis, is that transient changes in tonic inhibition during LTP induction may also modulate the outcome of LTP induced by TBS under different physiological conditions, since both synaptic and non-synaptic inhibition can hinder spike backpropagation and Ca^2+^ spikes in CA1 pyramidal cell dendrites, a mechanism believed to be required for TBS-induced LTP (Tsubokawa and Ross, 1996; Larson and Munkácsy, 2015; Müllner et al., 2015). In fact, developmental changes in tonic GABAergic inhibition at pyramidal cell dendrites have been associated with developmental changes in LTP expression (Groen et al., 2014), suggesting that such changes may contribute to the differences in the magnitude of potentiation induced by *mild* TBS at the three developmental stages tested in this study. In addition, induction of iLTD by TBS may influence the outcome of TBS-induced LTP at glutamatergic synapses by altering the GABAergic tone to pyramidal cell dendrites (Artinian and Lacaille, 2018; Udakis et al., 2020).

In this study, LTP induced by TBS was, as expected, dependent on endogenous activation of GABA_B_ receptors, as previously described, thus confirming an essential role of these receptors in CA1 LTP induction by TBS (Larson et al., 1986; Larson and Lynch, 1988). However, although blockade of GABA_B_ receptors with supramaximal concentrations of the antagonist CGP 55845 inhibited TBS-induced LTP in the CA1 area of the hippocampus, in line with previous studies (Lecouflet et al., 2021), it did not prevent the expression of TBS-induced LTP in this study. This suggests that additional GABA_B_-independent mechanisms contribute to the suppression of feed-forward inhibition to induce LTP by TBS. In addition, blockade of GAT1 transporters, that are predominantly expressed in presynaptic GABAergic boutons and to a lesser extent in astrocytes, enhanced LTP induced by *mild TBS* in juvenile rats, rebutting what was previously reported in mice using the same stimulation paradigm and intensity (Gong et al., 2009). GABA released to the synaptic cleft is subject to reuptake by high-affinity Na^+^- and Cl^-^-dependent GABA transporters (GATs). GAT-1 plays a crucial role in controlling synaptic GABA spillover and, consequently, modulation of both phasic and tonic GABAergic inhibition (Gong et al., 2009). In this paper we used submaximal inhibition of GAT-1 transporters to demonstrate their role in controlling the outcome of LTP induced by mild TBS in order to avoid non-synaptic effects of enhanced synaptic GABA through GABA spillover and to avoid non-specific effects of SKF89976A on GAT-3 transporters. In light of the results in this study, it is suggested that GAT-1 blockade, by enhancing synaptic GABAergic tone at synapses mediating disinhibition, might contribute to enhanced depolarization of pyramidal cell dendrites during TBS, leading to enhanced potentiation of glutamatergic transmission (Cunha-Reis et al., 2010, 2014; Francavilla et al., 2018; Cunha-Reis and Caulino-Rocha, 2020). In fact, recent evidence strongly suggests that disinhibition, can control CA1 synaptic plasticity, through regulation of long-term depression of inhibition (iLTD) and long-term potentiation of inhibition (iLTP) (Artinian and Lacaille, 2018; Udakis et al., 2020). Enhanced synaptic GABA by GAT-1 inhibition at feedforward inhibitory contacts might enhance GABA_B_ mediated suppression of inhibition, leading to increased potentiation by TBS. The observed inhibition of mild-TBS-induced LTP by GAT-1 inhibition in mice may otherwise reflect the enhanced feed-forward inhibition by enhanced GABAergic tone on pyramidal cell dendrites. The precise synaptic mechanisms of GAT-1 blockade-mediated changes in GABAergic disinhibition and inhibition in rats and mice could only be fully clarified by performing synaptic recordings in pyramidal cells and interneurons, including interneuron-selective interneurons. Altogether, this suggests that theta-patterned neuronal activity may be differently regulated in mice and in rats during exploration and spatial learning, the behavioral correlates of the *mild TBS* LTP-inducing paradigm characterized in this study. In fact, mice and rats behave very differently in exploratory tasks and its modulation by sensory cues and previous experience are very distinct (Tanaka et al., 2012; Thompson et al., 2018). Interestingly, Reyes-Garcia et al (2018) described that an enhancement of TBS-induced LTP by GABA_A_ receptor antagonists is not observed in hippocampal slices from a rat wildlife population (Reyes-Garcia et al., 2018). The enhancement in *mild* TBS-induced LTP expression caused by environmental enrichment (Malik and Chattarji, 2012) may involve changes in number and distribution of GABAergic synapses onto pyramidal cell dendrites (Foggetti et al., 2019). Furthermore, the enhancement of mild TBS-induced LTP caused by mismatch exploration training (Aidil-Carvalho et al., 2017) is also associated with a decrease in VIP-mediated endogenous disinhibition enhancing LTP (Cunha-Reis, 2020). Altogether, these observations and the fact that variations in LTP across dendritic subfields of hippocampus pyramidal cells reflect a differential distribution of GABAergic interneurons responsible for feed-forward inhibition, that in turn differentially affect the local dynamics of Ca^2+^ transients (Arai et al., 1994; Kanemoto et al., 2011; Francavilla et al., 2019), suggest that a dynamic modulation of GABAergic input to different CA1 pyramidal cell dendritic subfields mediates changes in *mild* TBS-induced LTP (Müllner et al., 2015).

In this study, the long-lasting potentiation of synaptic transmission caused by mild TBS in young adult rats was fully dependent on NMDA receptor activation and CaMKII activity (Fig. 7). This was likely a result from enhanced Ca^2+^ influx (Fig. 3. A-B), as previously described (Larson and Munkácsy, 2015), and does not suggest that additional endogenous channel activation is required to induce potentiation under these experimental conditions. Furthermore, as previously described by us and others (Cruz, 2010; Park et al., 2016, 2021), this potentiation was independent of PKA activity (Fig. 3.C). Even when reproducing experimental conditions and using specific inhibitors of PKA activity, as previously used in mice when observing a PKA-dependent expression of theta-burst LTP, this independence of PKA activity was confirmed (Fig. 4). This again suggests that TBS-induced LTP is differently regulated in rats and in mice, and that these differences should not be disregarded in the context of learning and memory tasks in the two species. This also implies that modulators acting through PKA-dependent mechanisms might have substantially different impact in the expression of TBS-induced LTP in mice and in rats, and that care should be taken in generalizing the interpretation of data from a single rodent species when considering substances acting through PKA-dependent transduction pathways. Interestingly, it has been described that PKA activity contributes to HFS-induced CA1 LTP in female but not in male Sprague-Dawley rats (Jain et al., 2019). Although elicited by a different stimulation pattern than the one used in this study these results corroborate the view that PKA activity is not universally required for LTP expression. In addition, recent studies describe that TBS-induced LTP induced by differently spaced sets of TBS patterns in the rat has different reliance on PKA activity (Park et al., 2016, 2021), with stronger spaced sets of TBS eliciting PKA-dependent processes. However, the physiological relevance of these different TBS intensities/patterns is still elusive. Finally, the expression of LTP induced by *mild* TBS was virtually independent of PKC activity. PKC has previously been reported to be involved in the expression of LTP induced by weak TBS stimulation 3×5 (100Hz) pulses in the CA1 area of mice hippocampal slices (Miura et al., 2002), but not in the rat (Hasegawa et al., 2014). These results suggest once more that regulation of LTP expression in the mouse and rat hippocampus is differently regulated by intracellular protein kinases.

The results in this study showed that *mild* TBS promotes phosphorylation of AMPA receptor GluA1 subunits at Ser_831_, a phosphorylation site targeted by both CaMKII and PKC, but not at Ser_845_, targeted by PKA. Phosphorylation of AMPA receptor GluA1 subunits by CaMKII, promotes enhancement of channel conductance and its recruitment of to the active zone (Derkach et al., 1999; Appleby et al., 2011). Although this has been argued not to be essential to the expression of hippocampal NMDA-dependent LTP (Henley and Wilkinson, 2016) or required for LTP stability (Henley and Wilkinson, 2016; Benke and Traynelis, 2019), the results in this study indicate than GluA1 phosphorylation at Ser_831_ occurs even upon *mild* TBS stimulation, suggesting that it might be required for expression of TBS-induced LTP. A very recent study corroborates these observations, by showing that CamKII activity is necessary for expression of CA1 hippocampal TBS-induced LTP (Park et al., 2021). Phosphorylation of GluA1 at Ser_845_, dependent on PKA activity and governing the insertion of calcium-permeable AMPA receptors to the synapse, was shown to be dependent on the TBS stimulation pattern and intensity used *in vitro* in the rat (Huang Yan You and Kandel, 1994; Park et al., 2016, 2021). Furthermore, since LTP induced by *moderate* TBS in mice hippocampal slices is PKA-dependent (Nguyen and Kandel, 1997), it is also dependent on the rodent species used. This again evidences a major variability in the role of GluA1 phosphorylation on *mild* to *moderate* TBS induced LTP in different rodent models. Furthermore, it is dependent on the use of maximal and submaximal TBS stimulation, hindering the establishment of an universal rule for the role of GluA1 phosphorylation in the induction, expression, and maintenance of TBS-induced LTP.

A role for Kv4.2 channels in regulation of hippocampal synaptic plasticity has long been debated in the literature (Kim and Hoffman, 2008). The function and properties of the Kv4.2 channel can be regulated by post-translational phosphorylation at several highly conserved sites. Kv4.2 can be phosphorylated by ERK and other kinases of the MAPK (mitogen-activated protein kinases) family at Thr_602_ and Thr_607_ residues, which leads to a decrease in I_A_, and internalization of the channel (Adams et al., 2000; Varga et al., 2000; Schrader et al., 2006). Yet kinases of the ERK family can also catalyze the phosphorylation of Ser_616_, which inhibits the channel’s activity (Hu et al., 2006). PKA phosphorylates the Kv4.2 channel, at Thr_38_ or Ser_552_. Although consequences of phosphorylation on Thr_38_ are unknown, phosphorylation of Ser_552_ causes internalization of the channel (Hammond et al., 2008). Finally, CaMKII can modulate Kv4.2 channel expression and upregulate neuronal I_A_ currents through phosphorylation at Ser_438_ (Varga et al., 2004).

One important observation in this study was that *mild* TBS stimulation enhances the phosphorylation of Kv4.2 channels at Ser_438_. As mentioned above, NMDA receptor activation during *mild* TBS mediates Ca^2+^ influx that activates CaMKII. This enhancement of CaMKII activity can be the cause for the enhanced Kv4.2 channel phosphorylation at Ser_438_ found in this work since this is a major target for CaMKII phosphorylation. This post-translational modification leads to an increase in the cell-surface expression of the channel (Varga et al., 2004), which can explain the mild increase of the channel levels also found in this work. Since the increase of the channel expression leads to an increase of I_A_ this likely does not contribute to expression of *mild* TBS-induced LTP since it promotes an increase in the NR2B/NR2A subunit ratio of the NMDA receptor (Jung et al., 2008, 2011). Rather it may be counteracting the effects of NMDA receptor activation to keep dendritic excitability under control. Furthermore, since Kv4.2 channels are differentially expressed in different dendritic segments in relation to the distance to the soma of pyramidal cells (Hoffman et al., 1997; Beck and Yaari, 2008) it is possible that these results do not reflect the modulation of the I_A_ in all dendritic segments. In addition, we did not find any evidence for phosphorylation of Kv4.2 channels at Thr_607_ and Thr_602_ phosphorylation sites targeted by ERK (extracellular signal-regulated kinases). Given the experimental design, mild TBS stimulation of a restricted group of synapses, and the low levels of Thr_607_ and Thr_602_ basal phosphorylation, it is possible that local changes in Kv4.2 phosphorylation by ERK may have been missed and could be detected by a more sensitive experimental approach. In fact, a physiological role for ERK in regulation of postsynaptic spiking following a strong TBS pattern has been described (Selcher et al., 2003) and could be due to regulation of dendritic Kv4.2 channels (Kim and Hoffman, 2008), although other molecular targets could not be excluded.

In summary, this paper shows that expression of LTP induced by *mild* TBS in the CA1 area of rat hippocampal slices involves several different signaling mechanisms than the ones currently accepted in the literature to be required for CA1 LTP expression (Fig. 7). One interesting study describes the mild vs strong TBS at perforant path (PP) synapses in the dentate gyrus (Jedlicka et al., 2015). Yet, the mild TBS used in that study is a lot stronger than the one used in this study, as it involves several episodes of TBS stimulation with larger interpulse interval and higher number of bursts, making a comparison quite difficult. Another important study reported different molecular mechanisms contributing to early LTP at CA3 to CA1 synapses with the ones required at PP-granule cells synapses using yet again a different pattern of stimulation between the different synapse sets (Cooke et al., 2006). As such, determining the more physiologically relevant stimulation patterns and intensities for each hippocampal subfield is imperative to the study of LTP, widely accepted as the required for learning and memory, represents still a challenge in current research. Nevertheless, it is our belief that *mild* TBS constitutes an ideal pattern and stimulation intensity to study the value of pharmacological tools aiming to modulate CA1 LTP expression and stability.

The results in this paper also raise several concerns on which is the best animal/rodent model to study the cellular mechanisms of LTP and the necessity to study both female and male LTP. In addition, this study evidences several differences in the endogenous regulation of TBS-induced LTP by GABA_A_ receptors and GAT-1 transporters which suggest that the GABAergic tone in the mouse and rat hippocampus during learning and memory may be quite different. This may in turn be the cause for the great differences in exploratory and learning behavior found between several mice and rat strains. Furthermore, we demonstrate that a few protein kinases widely accepted to be essential to LTP expression in mice do not significantly contribute to LTP expression induced by *mild* to *moderate TBS* in the rat, suggesting that they are more likely involved in regulatory mechanisms or activated by endogenous modulators of LTP in well-defined physiological conditions. We also show that GluA1 phosphorylation at Ser_831_ is the only GluA1 post-translational modification that is essential for expression of *mild TBS* induced LTP in the CA1 area of the rat hippocampus suggesting other phosphorylation sites in GluA1 subunits may be instead targets of endogenous modulators of synaptic plasticity. Finally, we show that Kv4.2 is more involved in suppressing or controlling firing than contributing to the expression of LTP induced by *mild TBS*. Nevertheless, its multiple phosphorylation sites suggest this channel is a major target for the regulation of LTP by endogenous modulators and several intracellular kinases. This may be very relevant for stronger stimulation intensities and patterns.

## Conclusion

In conclusion, this paper highlights several differences in the essential mechanisms involved in mild TBS-induced LTP in the rat hippocampal CA1 when compared to other stimulation patterns or animal models. This supports the need to establish the best animal model as well as standardized methodologies, stimulation patterns and intensities to induce LTP by TBS to determine which are the most suitable and relevant to study the action of pharmacological tools aiming to improve synaptic plasticity and ultimately memory.

## Acknowledgements

We acknowledge Prof. Alexandre de Mendonça for helpful discussions on TBS stimulation protocols and the Institute of Physiology, FMUL, for animal housing facilities.

## Fellowships

SFRH/BPD/34661/2007 and SFRH/BPD/81358/2011 to DCR and

## Researcher contract

Norma Transitória - DL57/2016/CP1479/CT0044 to DCR. Funding sources made no contribution to the writing, research plan and decision to publish this paper.

## Data accessibility statement

The data that support the findings of this study are available on request from the corresponding author. The data are not publicly available due to privacy or ethical restrictions.

## Ethics statement

The work in this project was approved by the Ethics committee of the Faculty of Medicine, University of Lisbon (Comissão de ética para a saúde do CHLN/FMUL). Approval was given in writing based on a detailed procedure report by the authors.A version of this paper was published as a preprint in BioRXiv doi: https://doi.org/10.1101/2021.04.19.440440.

## Abreviations

aCSF: artificial cerebrospinal fluid
AMPAR: α-amino-3-hydroxy-5-methyl-4-isoxazolepropionic acid receptor
AP-5: (2R)-amino-5-phosphonopentanoate
ATP: adenosine triphosphate
CA1: *Cornu Ammonis* 1
CaCl_2_: Calcium chloride
CaMKII: calmodulin-dependent protein kinase II
CO_2_: carbon dioxide
EDTA: Ethylenediamine tetraacetic acid
EEG: Electroencephalography
ERK: extracellular signal-regulated kinases
fEPSP: field excitatory post-synaptic potential
GABA: Gamma Aminobutyric Acid
GAT1: GABA transporter 1
GATs: GABA transporters
GluA1: AMPA subunit glutamate receptor 1
HEPES: 4-(2-hydroxyethyl)-1-piperazineethanesulfonic acid
HFS: high-frequency stimulation
HRP: horseradish peroxidase
I_A_: fast activating and deactivating A-current
KCl: potassium chloride
Kv4.2: voltage-gated potassium channel subunit 4.2
LTD: long-term depression
LTP: long-term potentiation
MgSO_4_: magnesium sulfate
Na_3_VO_4_: sodium orthovanadate
NaCl: sodium chloride
NaF: sodium floride
NaH_2_PO_4_: sodium phosphate monobasic monohydrate
NaHCO_3_: sodium bicarbonate
NMDAR: N-methyl-D-aspartate receptor
PKA: protein kinase A
PKC: protein kinase C
PMSF: phenylmethylsulfonyl fluoride
PPF: paired-pulse facilitation
PVDF: polyvinylidene fluoride
SDS-PAGE: sodium dodecyl sulphate polyacrylamide gel electrophoresis
VIP: vasoactive intestinal peptide

